# Field potential Imaging in human iPSC- derived Cardiomyocytes using UHD-CMOS-MEA

**DOI:** 10.1101/2025.03.31.646249

**Authors:** Naoki Matsuda, Nami Nagafuku, Kazuki Matsuda, Yuto Ishibashi, Tomohiko Taniguchi, Yusaku Matsushita, Norimasa Miyamoto, Takashi Yoshinaga, Ikuro Suzuki

**Affiliations:** Department of Electronics, Graduate School of Engineering, Tohoku Institute of Technology, 35-1 Yagiyama Kasumicho, Taihaku-ku, Sendai, Miyagi, 982-8577, Japan; Advanced Biosignal Safety Assessment, Biopharmaceutical Assessments Unit, Eisai Co., Ltd., 5-1-3 Tokodai, Tsukuba, Ibaraki 300-2635, Japan

## Abstract

Evaluation using beat propagation analysis of human iPS cardiomyocytes is an effective approach for assessing human cardiac safety in drug development. However, current applications are primarily focused on detecting QT prolongation and arrhythmia risk, while its ability to comprehensively detect cardiotoxicity remains insufficient. Additionally, predicting the mechanism of action, which is crucial in drug development, remains challenging. In this study, we employed field potential imaging (FPI) using an ultra-high-density (UHD) CMOS microelectrode array (MEA) comprising 236,880 electrodes with high spatiotemporal resolution, capable of recording the activity of a monolayer of cardiomyocytes with tens of electrodes per cell. This method enabled the establishment of novel electrophysiological endpoints, including the number of excitation origins, fluctuations in origin positions, conduction velocity, and propagation area. Pharmacological characterization revealed drug-specific effects: isoproterenol increased excitation origins, mexiletine reduced conduction velocity, and E-4031 decreased propagation area while inducing early afterdepolarizations. Multivariate analysis of 13 compounds across 17 electrophysiological endpoints distinguished conduction velocity and propagation patterns based on their mechanisms of action. Additionally, 0.1 μM doxorubicin exposure for 24 hours significantly reduced conduction velocity and propagation area, allowing early detection of chronic cardiotoxicity. These findings suggest that UHD-CMOS-MEA-based FPI enhances cardiotoxicity detection at low concentrations and precisely characterizes ion channel activity across different drug concentrations. The integration of novel electrophysiological endpoints derived from UHD-CMOS-MEA-based FPI, including excitation origin analysis, conduction velocity, and propagation area, along with multivariate analysis, is anticipated to establish a next-generation in vitro platform for comprehensive cardiotoxicity risk assessment and mechanism-based prediction of drug candidates.

## Introduction

Cardiotoxicity is one of the most serious adverse effects in drug development, posing a significant threat to patient safety^1, 2, 3, 4^. Therefore, its evaluation is essential to ensure the safety of new drugs. In particular, drug-induced QT prolongation and Torsades de Pointes (TdP) arrhythmia are widely recognized as critical risks, often leading to the termination of clinical trials or withdrawal of marketed drugs^5^.

Traditionally, cardiotoxicity assessment for drug candidates has been based on hERG channel inhibition assays and animal models. The ICH S7B guideline has played a pivotal role in standardizing the evaluation of QT prolongation risk due to hERG channel inhibition, establishing a foundation for cardiotoxicity assessment^6, 7^. In recent years, the Comprehensive in vitro Proarrhythmic Assay (CiPA) initiative has driven a paradigm shift in this field^8, 9, 10^. The development of the ICH E14/S7B Q&A document has facilitated the transition from a reliance on hERG channel inhibition alone to a more integrated assessment of proarrhythmic risk. This approach incorporates multiple ion channel evaluations, human-derived cardiomyocyte assays, and in silico models, allowing for a more comprehensive assessment of not only QT prolongation but also the actual risk of arrhythmia induction^8, 11^. Consequently, this methodology has been adopted by regulatory agencies, including the FDA, for drug approval application.

However, several challenges remain within the frameworks of ICH S7B and ICH E14/S7B Q&A. The ICH S7B guideline primarily focuses on the evaluation of hERG channel inhibition and QT prolongation but does not fully address a comprehensive analysis of mechanisms involving other ion channels, such as Na^+^ and Ca^2+^ channels^8, 9^. On the other hand, while the ICH E14/S7B Q&A has enhanced the accuracy of TdP risk prediction by integrating data from multiple ion channels, fully elucidating the hierarchy of effects and the impact of interactions remains challenging for compounds that act on multiple ion channels simultaneously. For example, in vitro data may not always correlate with clinical outcomes for compounds such as verapamil and bepridil, which inhibit hERG channels while also modulating L-type Ca^2+^ channels (ICaL) and Na^+^ channels (INa)^12^. Furthermore, CiPA primarily focuses on the acute assessment of proarrhythmic risk and does not comprehensively evaluate chronic toxicity or the detailed conduction of electrical activity in cardiomyocytes.

Against this backdrop, the utilization of human induced pluripotent stem cell-derived cardiomyocytes (hiPSC-CMs) as an in vitro evaluation system has been expanding^13^. hiPSC-CMs have the potential to more accurately detect toxicity risks that are difficult to capture in animal models, as they better recapitulate human-specific electrophysiological properties^14, 15, 16, 17, 18^. From a regulatory perspective, the assessment of chronic toxicity is increasingly required. However, hiPSC-CMs remain immature compared to adult cardiomyocytes, exhibiting differences in ion channel expression profiles and intracellular Ca²⁺ handling mechanisms. Consequently, the evaluation results obtained using hiPSC-CMs may not always correlate with actual clinical toxicity^14^. Despite their utility as a valuable evaluation system, the reduced predictive accuracy of toxicity due to their immaturity remains a major challenge. Therefore, optimizing culture conditions to enhance maturation and developing technologies that enable higher-resolution electrophysiological assessments are essential for improving the reliability of hiPSC-CM-based cardiotoxicity evaluations.

Following the development of culture techniques, enhancing cellular maturation is crucial. However, improving measurement technologies to enable the detection of previously undetectable cardiotoxicity and to predict mechanisms of action is even more important and provides a more practical solution. In recent years, various in vitro evaluation systems have been developed and utilized, including impedance measurements for real-time assessment of cellular morphological changes and contractile force, Ca²⁺ transient measurements for visualizing intracellular Ca²⁺ dynamics, and high-content imaging for capturing cellular morphology and contraction^19, 20, 21, 22, 23, 24^.

Impedance measurements offer the advantage of high sensitivity in detecting subtle morphological changes and interactions with the extracellular matrix. Meanwhile, Ca^²⁺^ transient measurements enable real-time monitoring of excitation-contraction coupling. However, each of these methods has limitations in directly evaluating membrane potential and spatial propagation of field potentials, making them insufficient for comprehensively reflecting complex interactions among multiple ion channels. In contrast, multi-electrode arrays (MEAs) allow for non-invasive, multi-site membrane potential recordings, facilitating the assessment of conduction velocity and electrical remodeling in cardiomyocytes, thereby enabling a more comprehensive cardiotoxicity analysis^12, 13, 15, 25, 26, 27, 28, 29, 30, 31^. However, traditional MEAs still have limitations in electrode density, as field potential duration (FPD) measurements from individual electrodes have been the primary method of evaluation, restricting the amount of information obtained. The integration of large-area, high-density CMOS technology presents a promising approach for uncovering previously undetectable functional characteristics of cardiomyocyte populations^32, 33, 34^.

In this study, we propose a novel approach, Field Potential Imaging (FPI) in hiPSC-derived Cardiomyocytes, utilizing an ultra-high-density (UHD) CMOS-MEA with 236,880 electrodes to address the aforementioned challenges. This method enables the visualization of fine-scale action potential propagation within cardiomyocyte populations, allowing for a multi-faceted and high-precision evaluation of cardiotoxicity in compounds targeting different mechanisms of action. Specifically, we comprehensively analyze novel electrophysiological endpoints, including the number of excitation origins, propagation area and velocity, beat intervals, and field potential amplitude. We demonstrate that this approach is applicable not only for assessing acute toxicity but also for detecting chronic toxicity risks, such as those associated with the anticancer drug doxorubicin. Furthermore, by leveraging the high spatial and temporal resolution of the CMOS-MEA, we validate its capability for more reliable toxicity assessment, including the estimation of mechanisms of action, even for compounds such as bepridil, which has been classified as low-risk under CiPA. This study provides a new platform for improving the accuracy of cardiotoxicity evaluation in drug development and for capturing complex interactions among multiple ion channels.

## Results

### Results 1. Field Potential Imaging of Human iPSC-derived Cardiomyocytes Using 236,880-electrode UHD-CMOS-MEA Measurements

Human iPSC-derived cardiomyocytes were cultured on a CMOS-MEA chip for 7–10 days in vitro (DIV), and field potential imaging was performed to capture the activity of all cardiomyocytes. Phase-contrast microscopic observation confirmed that the cardiomyocytes adhered uniformly to the electrode area and that a single cell spanned multiple electrodes of the CMOS-MEA (Fig. 1A), demonstrating the high-resolution capability to simultaneously record field potentials from multiple electrodes. Extracellular potential waveforms of spontaneously beating cardiomyocytes, recorded at a sampling frequency of 2 kHz, showed high-density and wide-range propagation of beating signals across the cell population (Fig. 1B). Figure 1B-a illustrates the waveform acquisition area within the cell seeding region. Figure 1B-b presents the raw waveform obtained at the red dot in 1B-a, clearly indicating the propagation of beats (peak voltage: 1.98 ± 0.20 mV, n = 9 electrodes). Furthermore, time-lapse analysis of potential heatmaps from each electrode at every sampling point revealed that a single beat propagated over 13 ms (Fig. 1C, Supplementary movie1).

**Figure 1.**
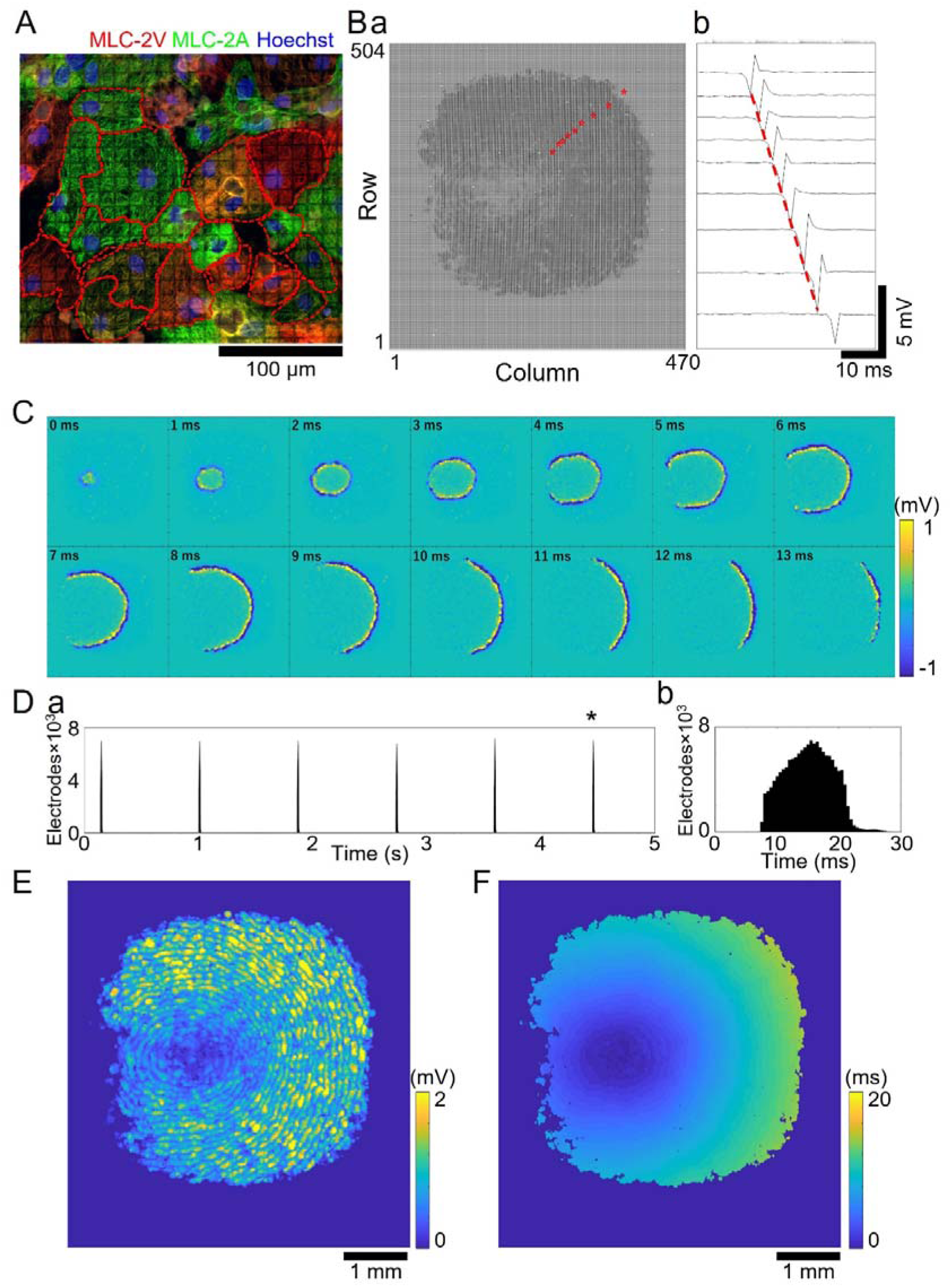
Field Potential Imaging in human iPSC-derived cardiomyocytes. (A) Immunostaining images. Red: Myosin Light Chain 2 Ventricular (MLC-2V); Green: Myosin Light Chain 2 Atrial (MLC-2A); Blue: Hoechst 33258. (B) (a) Extracellular potential waveforms recorded from 236,880 electrodes. (b) Extracellular potential waveform from the electrode marked with a star. (C) Time-lapse of the potential heatmap for each sampling point across all electrodes during a single beat. (D) (a) Histogram of the number of active electrodes over 5 seconds (bin size = 0.5 ms). (b) Enlarged view of a single beat marked with an asterisk in (a). (E) Heatmap of the maximum voltage amplitude during a single beat. (F) Delay map of voltage peak timing during a single beat.

Using extracellular potential waveforms, active electrodes were detected based on potential thresholds, and a histogram of active electrode counts was generated with a bin size of 0.5 ms (Fig. 1D-a). A single beat was detected across 115,892 ± 50 electrodes (n = 6 beats, Fig. 1D-b). By identifying the beating intervals and the peak number of active electrodes from the histogram, the beat count and interbeat intervals (IBI) could be evaluated (Fig. 1D). The results showed a stable and regular beating pattern, with an average beating rate of 57.8 ± 1.7 bpm and an average IBI of 1078.6 ± 29.8 ms (n = 54 wells, Fig. 1D). Additionally, heatmaps of peak voltage at each electrode and delay maps based on the peak timing of the voltage were created. These maps enabled detailed visualization of spatial propagation dynamics, including the initiation site, propagation pathway, and propagation area of a single beat (Fig. 1E). These findings demonstrate that field potential imaging of human iPSC-derived cardiomyocytes is feasible using UHD-CMOS-MEA. Moreover, the high spatial and temporal resolution of this system allows for comprehensive analysis of field potential propagation.

### Results 2. Establishment of Novel Endpoints for Beat Propagation and Detection of Drug Responses

The use of CMOS-MEA enables high spatial resolution measurements, allowing not only the assessment of beating frequency but also the precise identification of the initiation site of each beat. By creating heatmaps from the peak voltage and peak timing of field potentials recorded via CMOS-MEA, we identified the initiation sites of individual beats and established the number and positional variation of initiation sites as novel analytical parameters. Figure 2A shows histograms of the number of active electrodes before and after the treatment of isoproterenol, a β1-adrenergic receptor agonist. Although no significant changes were observed in the number of active electrodes following isoproterenol treatment (Before: 149,149.4 ± 3,090.5 electrodes, n = 71 beats; 10 µM: 149,185.2 ± 2,882.6 electrodes, n = 114 beats), an increase in beating frequency was observed (Before: 35.5 bpm, 30nM: 57.0 bpm), demonstrating the positive chronotropic effect of isoproterenol in MEA recordings. Figure 2B presents heatmaps of beat propagation time. Before isoproterenol treatment, beats propagated solely from the initiation site in the upper left region toward the lower right. However, after treatment, multiple initiation sites appeared in the lower right region in addition to the initial upper left site, indicating the induction of ectopic firing by isoproterenol, as also observed in MEA recordings (Supplementary movie 2). A plot of all initiation sites detected over a 120-second period confirmed an increase in initiation site count following isoproterenol treatment (Figure 2C). The number of initiation sites per beat significantly increased to 1.51 ± 0.02 and 1.61 ± 0.02 at isoproterenol concentrations of 10 nM and 30 nM, respectively, compared to pre-treatment values (n = 6 wells, at least 626 beats; Fig. 2D).

**Figure 2.**
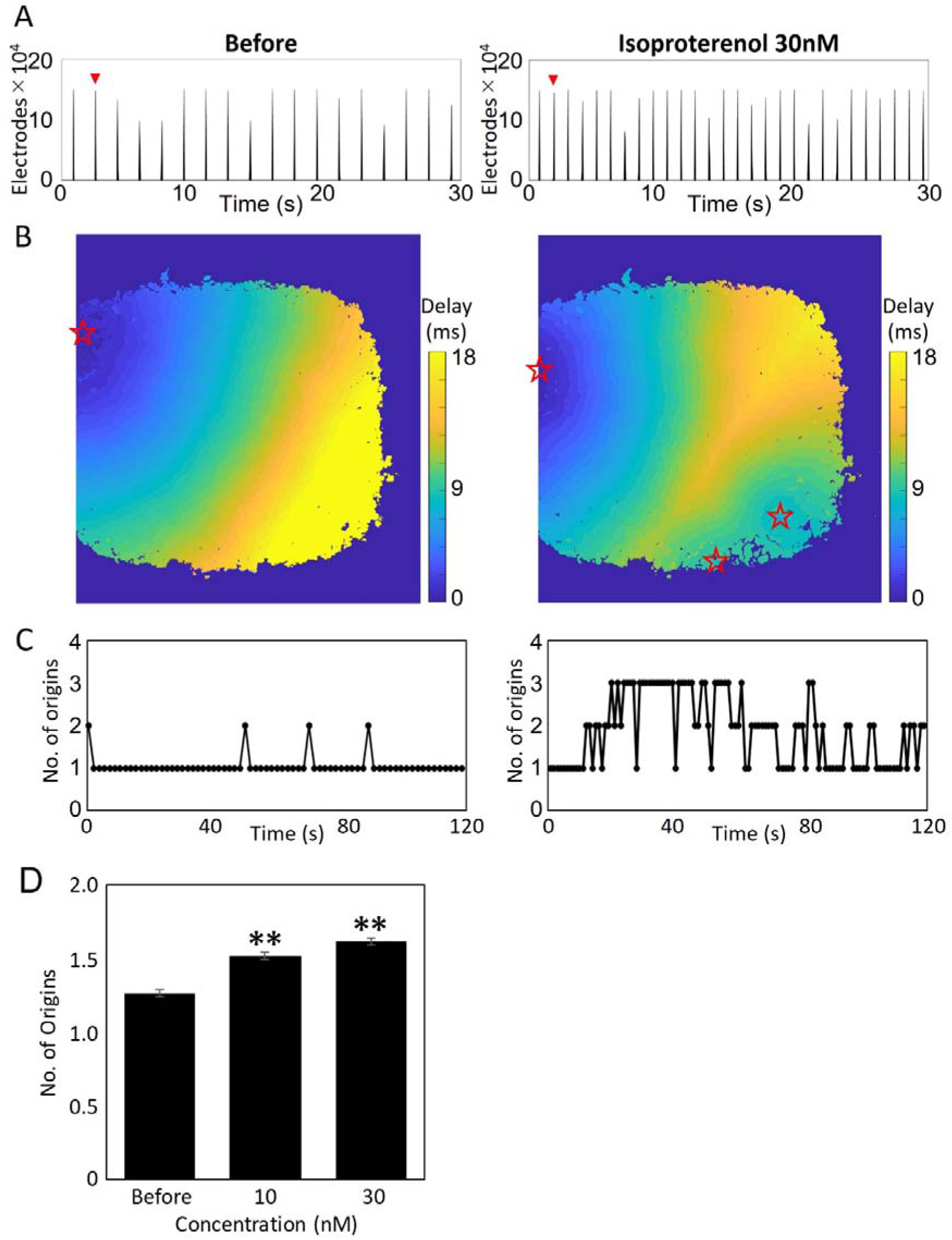
Increase in the Number of Excitation Origins Induced by Isoproterenol. (A)Histogram of the number of active electrodes over 30 seconds before (left) and after the treatment of 30 nM Isoproterenol (right). (B) Delay map of voltage peak timing for a single beat, where stars indicate excitation origins. (C) Plot of the number of excitation origins detected over a 2-minute period. (D) Concentration-dependent increase in the average number of excitation origins. (n = 6 wells, ** p < 0.01 vs. before)

CMOS-MEA-based measurements possess not only high spatial resolution but also high temporal resolution, enabling precise calculation of conduction velocity during beat propagation. Specifically, using the initiation site as a reference, a scatter plot was generated to illustrate the conduction distance and time for each electrode, and the overall conduction velocity was determined from the slope of the regression line. Additionally, the conduction velocity at each electrode was calculated based on the sum of the distances between the nearest electrode that fired 0.5 ms earlier and the nearest electrode that fired 0.5 ms later. A heatmap was then generated to visualize local variations in conduction velocity within a single beat. Figure 3A presents histograms of the number of active electrodes before and after the treatment of mexiletine, a sodium channel inhibitor. Mexiletine treatment resulted in a decrease in the number of active electrodes (Before: 115437.9 ± 582.1 electrodes, 10 µM: 93595.62 ± 11323.1 electrodes, n ≥ 63 beats) and a reduction in beating frequency (Before: 64.5 bpm, 10 µM: 31.5 bpm). Figure 3B shows the delay map of beat propagation time and the heatmap of conduction velocity before and after mexiletine treatment. Although beats continued to propagate throughout the entire region post-treatment, the proportion of regions with conduction velocities exceeding 0.3 m/s decreased from 4.9% to 0.6%, indicating an overall slowing of conduction velocity (Supplementary movie 3). An analysis of conduction velocity across individual beats revealed that, while conduction velocity remained consistent across beats before treatment, significant beat-to-beat variations emerged post-treatment, with some beats exhibiting markedly slower conduction velocities (Figure 3C). The mean conduction velocity showed a concentration-dependent decrease from 0.147 ± 0.004 m/s before mexiletine treatment to 0.109 ± 0.009 m/s at 10 µM, with a statistically significant difference observed (n = 6 wells; Fig. 3D).

**Figure 3.**
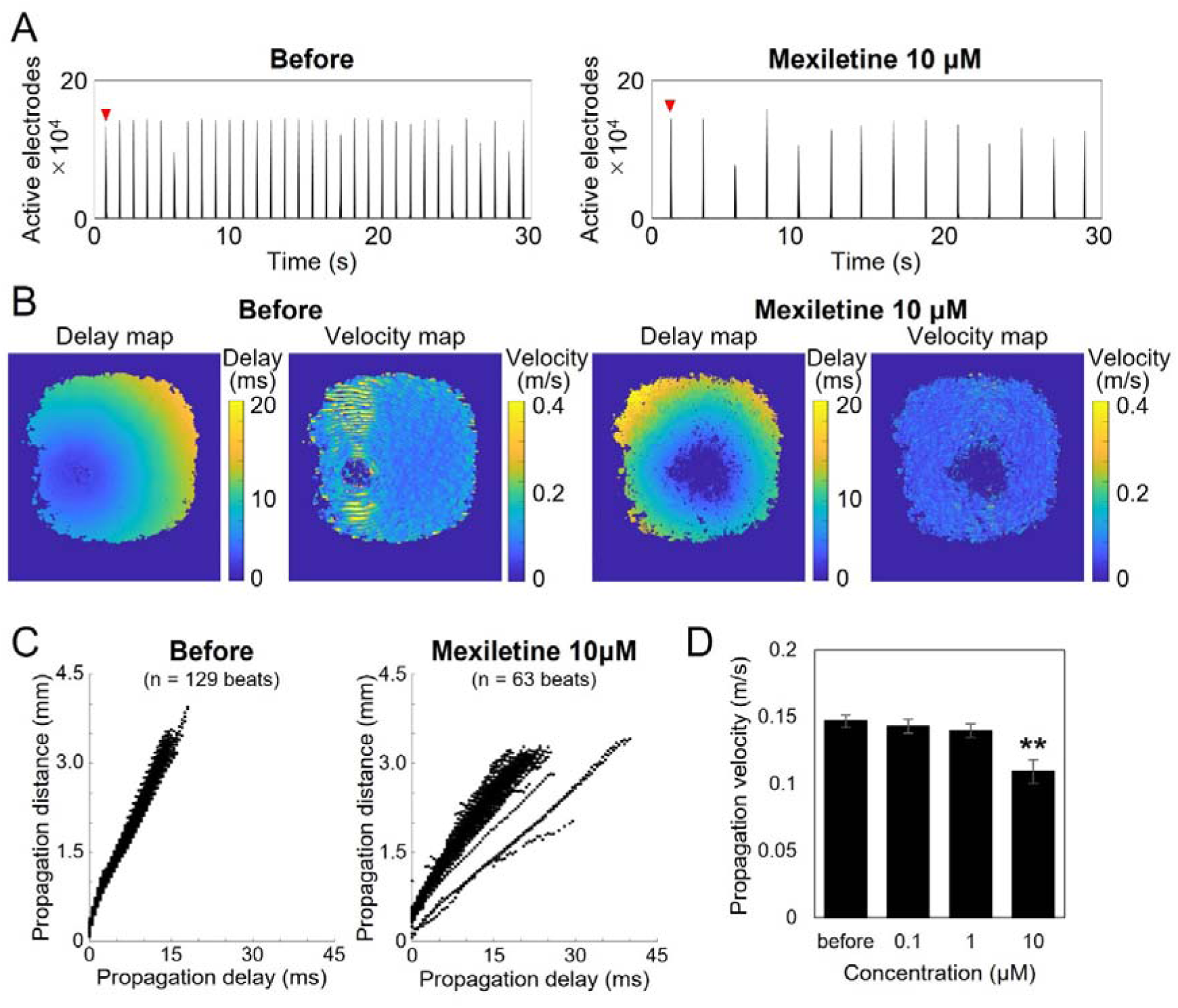
Decrease in Propagation Velocity Induced by Mexiletine. (A) Histogram of the number of active electrodes over 30 seconds before (left) and after the treatment of 10 μM Mexiletine (right). (B) Delay map of voltage peak timing and propagation velocity heatmap for a single beat. (C) Plot of propagation time and propagation distance detected over a 2-minute period. (D) Concentration-dependent decrease in the average propagation velocity. (n = 6 wells, p < 0.01 vs. before)

Furthermore, CMOS-MEA measurements with high spatiotemporal resolution enable the precise detection of the entire propagation area of each beat without omission, allowing for the accurate calculation of the propagation area. Figure 4A presents histograms of the number of active electrodes before and after the treatment of E-4031, a hERG potassium channel inhibitor. E-4031 treatment did not significantly alter beating frequency at 100 nM (Before: 88 bpm, 100 nM: 82 bpm); however, it decreased the number of active electrodes (Before: 166785.2 ± 4264.5 electrodes, 100 nM: 43675.2 ± 42039.4 electrodes)—representing a reduction in propagation area—while increasing variability. This suggests that arrhythmia-like responses induced by E-4031 were also detected via MEA measurements. Figure 4A-b illustrates the heatmap representing the propagation area. Before the treatment of 100 nM E-4031, beats were observed across the entire recording area. However, post-treatment, beats frequently occurred only in localized regions, with the specific regions varying across beats. The mean propagation area decreased in a concentration-dependent manner from 17.98 ± 1.78 mm² before E-4031 treatment to 6.51 ± 1.51 mm² at 100 nM, with a statistically significant difference observed. Additionally, the coefficient of variation (CV) of the propagation area increased in a concentration-dependent manner from 0.029 ± 0.004 before E-4031 treatment, reaching statistical significance at 30 nM with a CV of 0.400 ± 0.182 (n = 5 wells; Fig. 4B).

**Figure 4.**
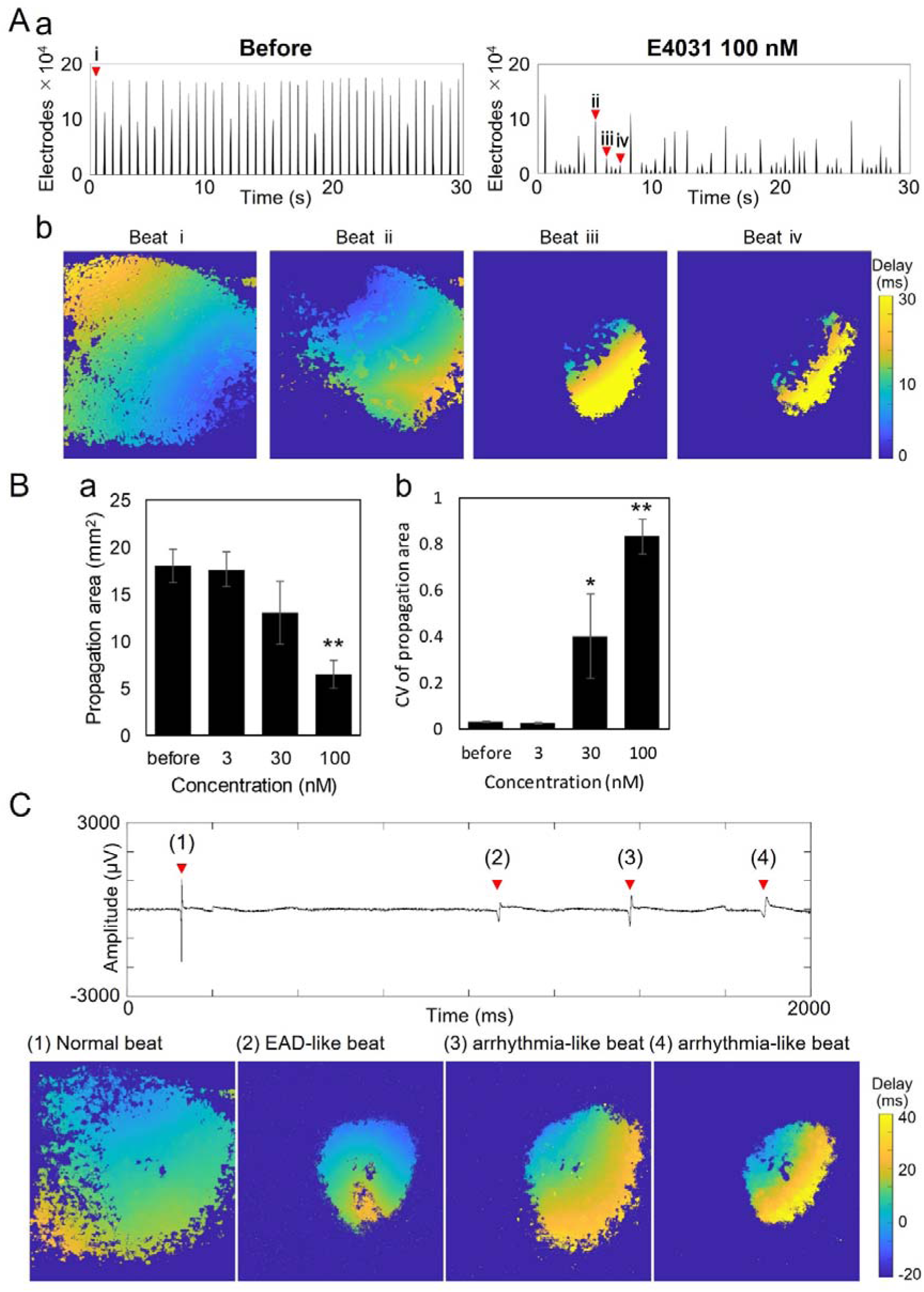
Decrease in Propagation Area and Detection of EAD-like Waveforms Induced by E-4031. (A) (a) Histogram of the number of active electrodes over 30 seconds before (left) and after the treatment of 100 nM E-4031 (right). (A) (b) Heatmap of peak timing for each beat indicated by arrows in (a). (B) (a) Concentration-dependent decrease in the average propagation area. (B) (b) Concentration-dependent increase in the coefficient of variation of propagation area (n = 6 wells). p < 0.01 vs. before. (C) (a) Field potential waveforms recorded after the treatment of 100 nM E-4031. (C) (b) Heatmap of peak timing for each beat indicated by arrows in (a). (1) Beats spreading across the entire recording area. (2) Locally confined beats with EAD-like waveforms. (3) Locally confined arrhythmic beats. (4) Locally confined arrhythmic beats with prolonged propagation time.

Additionally, at 30 nM E-4031, partial beats were observed immediately following widely propagated beats, with the extracellular field potential waveforms exhibiting early afterdepolarization (EAD)-like patterns similar to those clinically observed in arrhythmias (Fig. 4C). Figure 4C-a displays the extracellular field potential waveforms recorded from electrodes where partial beats were detected, while Figure 4C-b presents heatmaps of propagation times corresponding to the four red arrows in the waveforms. Following the widely propagated beat (1), a localized beat with prolonged propagation time was observed after 930 ms, followed by frequent occurrences of partial beats within the subsequent 400 ms (Fig. 4C, Supplementary movie 4). These newly identified beat parameters, evaluated through field potential imaging of human iPSC-derived cardiomyocytes, successfully distinguished differences in mechanisms of action.

### Results 3. Detection of Acute Cardiotoxicity and Classification of Mechanisms of Action Based on Beat Parameters

To validate the assessment of cardiotoxicity induced by acute drug treatment using CMOS-MEA measurements, we conducted FPI analysis on 11 compounds, including nine cardiotoxic positive compounds—some of which, such as Bepridil that conventional MEA could not detect—and two negative compounds. A total of 17 beat-related parameters were calculated (Table 1). Figure 5 presents a heatmap of the parameters for each compound. The negative compounds, aspirin and amoxicillin, showed no significant differences in any parameter. The selective hERG channel inhibitor E-4031 significantly reduced beat frequency, field potential amplitude, propagation area, conduction velocity, and the variability of origin locations. Additionally, localized beats with weak amplitudes, resembling early afterdepolarization (EAD), emerged, leading to increased variability in field potential amplitude and interbeat intervals (IBIs). Similarly, Disopyramide, which also affects the hERG channel, significantly reduced beat frequency, field potential amplitude, propagation area, conduction velocity, and origin variability. However, due to the decrease in beat frequency, the IBI was prolonged. The sodium channel blocker Mexiletine significantly reduced beat frequency, field potential amplitude, propagation area, and conduction velocity, while increasing the IBI interval. However, it did not alter the variability in IBIs, propagation area, or conduction velocity, with only field potential amplitude variability increasing. The IKs channel blocker Azimilide led to a decrease in beat frequency and an increase in IBI, along with a significant increase in IBI variability. The β-adrenergic receptor agonist Isoproterenol increased beat frequency, which was accompanied by a decrease in the variability of IBI and propagation area. Diltiazem, which acts on both CaV and hERG channels, increased beat frequency and decreased the IBI. Similarly, Verapamil increased beat frequency and decreased the IBI, but additionally increased field potential amplitude and decreased the distance of origins. These responses differed from those of E-4031 and Disopyramide, which selectively inhibit hERG channels. Quinidine and Bepridil, which act on CaV, NaV, and hERG channels, exhibited distinct responses. Quinidine reduced beat frequency, field potential amplitude, and propagation area, while increasing IBI and propagation area variability. Notably, its parameter changes resembled those observed with hERG channel inhibitors. Bepridil, on the other hand, decreased the variability of IBIs, field potential amplitude, and conduction velocity, demonstrating a composite effect distinct from any single-channel blocker. Table 2 summarizes the parameters that exhibited significant changes for each compound.

**Table 1.** Description of Analytical Parameters. A list of 17 analytical parameters along with detailed descriptions of the respective computational processes involved in their derivation.

| Analytical parameter | Description |
| --- | --- |
| Beat rate (BPM) | Number of peaks in the histogram of the active electrodes |
| Inter beat interbal (IBI, ms) | Average peak interval of histogram of active electrodes |
| CV of IBI | Coefficient of variation of IBI |
| STV of IBI | Short-Term Variance of IBI |
| Amplitude (mV) | Maximum absolute value of the amplitude of the action potential |
| CV of Amplitude | Coefficient of variation of Amplitude |
| STV of Amplitude | Short-Term Variance of Amplitude |
| Area (mm<sup>2</sup>) | Active electrodes area |
| CV of Area | Coefficient of variation of Area |
| STV of Area | Short-Term Variance of Area |
| Velocity | Average local conduction velocity calculated at each electrode location |
| CV of Velocity | Coefficient of variation of Velocity |
| STV of Velocity | Short-Term Variance of Velocity |
| No. of origins | Average number of origins per beat |
| Distance of origins | Average distance of change in the origins between two beats |
| CV of distance of origins | Coefficient of variation of Distance of origins |
| STV of distance of origins | Short-Term Variance of Distance of origins |

**Figure 5.**
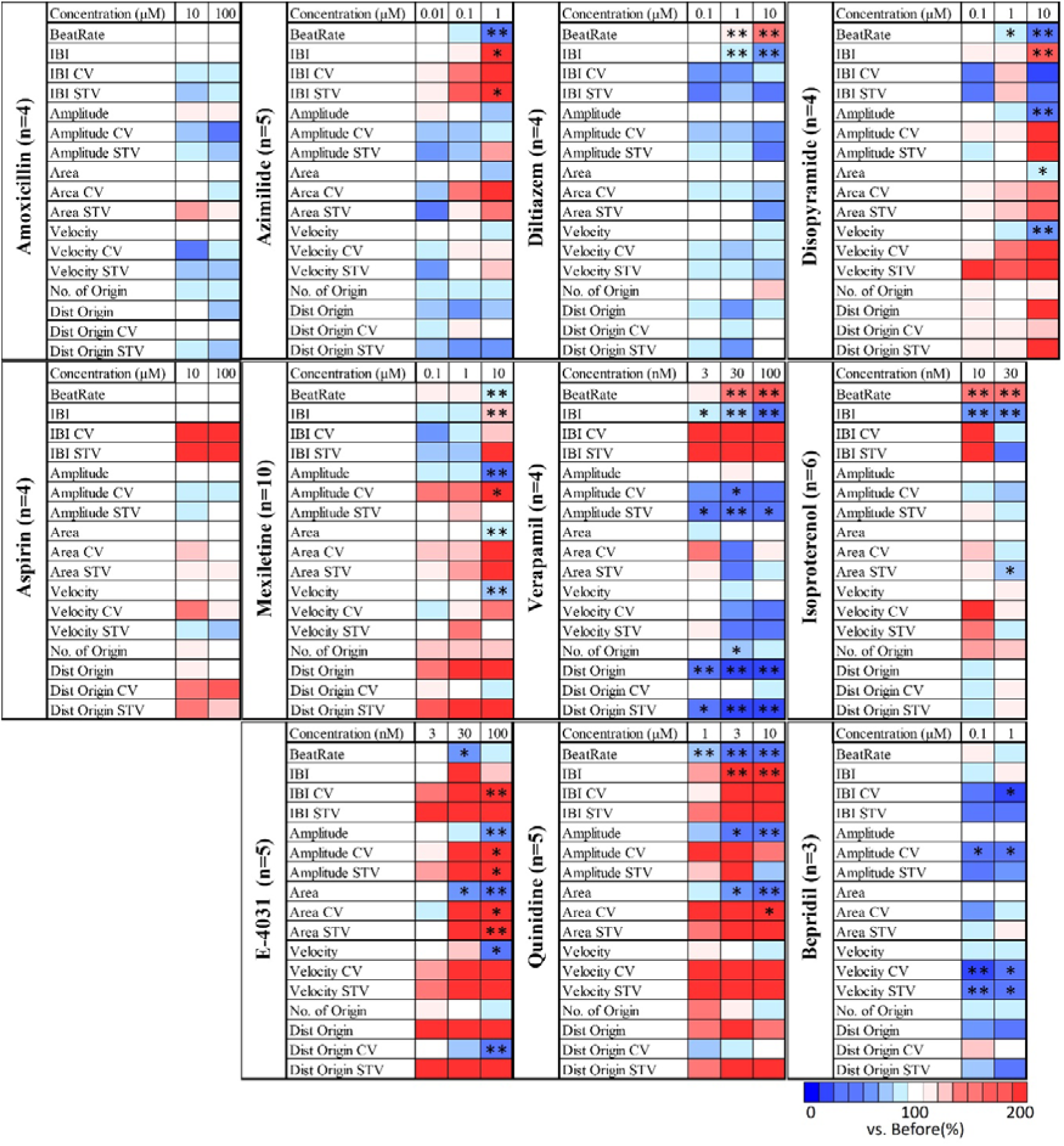
Heatmap of beating parameters of acute responses to cardiotoxicity-inducing compounds and negative controls in human iPSC-derived cardiomyocytes. Heatmap of beating parameters at each concentration, normalized to 100% based on pre-treatment values for each compound. One-way ANOVA followed by Dunnett’s test, *p < 0.05 vs. before, **p < 0.01 vs. before.

**Table 2.** Summary of quantitative compound effects on analytical parameters.

| **Compound**<br>**s** | **BR** | **IBI** | **IBI**<br>**CV** | **IBI**<br>**STV** | **Amp**<br>**.** | **Amp**<br>**. CV** | **Amp**<br>**. STV** | **Area** | **Area**<br>**CV** | **Area**<br>**STV** | **Velo**<br>**c.** | **Velo**<br>**c.**<br>**CV** | **Velo**<br>**c.**<br>**STV** | **No.**<br>**of**<br>**Orig**<br>**in** | **Dist**<br>**Orig**<br>**in** | **Dist**<br>**Orig**<br>**in**<br>**CV** | **Dist**<br>**Orig**<br>**in**<br>**STV** |
| --- | --- | --- | --- | --- | --- | --- | --- | --- | --- | --- | --- | --- | --- | --- | --- | --- | --- |
| **Diltiazem** | ↑ | ↓ | - | - | - | - | - | - | - | - | - | - | - | - | - | - | - |
| **Isoprotere**<br>**nol** | ↑ | ↓ | - | - | - | - | - | - | - | ↓ | - | - | - | - | - | - | - |
| **Verapamil** | ↑ | ↓ | - | - | - | ↓ | ↓ | - | - | - | - | - | - | ↓ | ↓ | - | ↓ |
| **Azimilide** | ↓ | ↑ | - | ↑ | - | - | - | - | - | - | - | - | - | - | - | - | - |
| **Disopyram**<br>**ide** | ↓ | ↑ | - | - | ↓ | - | - | ↓ | - | - | ↓ | - | - | - | - | - | - |
| **Mexiletine** | ↓ | ↑ | - | - | ↓ | ↑ | - | ↓ | - | - | ↓ | - | - | - | - | - | - |
| **E-4031** | ↓ | - | ↑ | - | ↓ | ↑ | ↑ | ↓ | ↑ | ↑ | ↓ | - | - | - | - | ↓ | - |
| **Quinidine** | ↓ | ↑ | - | - | ↓ | - | - | ↓ | ↑ | - | - | - | - | - | - | - | - |
| **Bepridil** | - | ↓ | ↓ | - | - | ↓ | - | - | - | - | - | ↓ | ↓ | - | - | - | - |
| **Amoxicillin** | - | - | - | - | - | - | - | - | - | - | - | - | - | - | - | - | - |
| **Aspirin** | - | - | - | - | - | - | - | - | - | - | - | - | - | - | - | - | - |

BR, Beat rate; Amp., Amplitude; Veloc., Velocity;

As described above, each compound exhibited characteristic changes in multiple parameters, such as beat frequency, field potential amplitude, propagation area, and conduction velocity, depending on its mechanism of action. However, it is challenging to comprehensively compare the differences and similarities between compounds using a single parameter. Therefore, in this study, we conducted principal component analysis (PCA) using four of the 17 extracted beat-related parameters—Beat Rate, IBI, Area, and Velocity STV—to comprehensively evaluate cardiotoxicity (Figure 6A). Table 3 presents the parameters used for PCA and their loading values (cumulative contribution rate of PC1 and PC2: 82.8%). PCA results showed that negative compounds remained near the origin, whereas positive compounds moved away from the origin in a concentration-dependent manner, each following a characteristic trajectory. Next, we established a toxicity threshold on the PCA plot based on the standard deviation (SD) range of negative compounds. Specifically, we defined the risk levels as follows: “within the SD range = low risk,” “between the SD and 2SD lines = medium risk,” and “outside the 2SD line = high risk” (Figure 6B). The PCA-based toxicity assessments were then compared with the IC_50_ and Cmax concentrations for each compound and ion channel (Table 4). The risk levels were color-coded as blue for low risk, pink for medium risk, and red for high risk, with star symbols indicating the IC_50_ and Cmax concentrations for each channel. As a result, all negative compounds were classified as low risk at all tested concentrations, while all positive compounds reached high-risk levels at certain concentrations. Importantly, this toxicity classification method was effective regardless of the differences in mechanism of action. For example, Diltiazem and Disopyramide were classified as high risk at concentrations exceeding their Cmax, whereas Azimilide, Bepridil, E-4031, and Quinidine reached the high-risk zone at concentrations near their IC_50_ for at least one channel. On the other hand, Mexiletine and Verapamil exhibited high risk at concentrations lower than their Cmax, demonstrating compound-specific differences in the concentration-dependent manifestation of toxicity. To investigate whether the PCA plots could distinguish different mechanisms of action, we categorized the obtained plots into six groups: Negative, CaV channel inhibitors, hERG channel inhibitors, hERG + CaV inhibitors, hERG + NaV inhibitors, and IKs inhibitors. We then performed statistical analysis using MANOVA (Figure 6C, Table 5). The results indicated that the Negative group, the IKs group and the hERG + NaV group showed significant differences from all other groups, suggesting that they could be distinctly separated based on their mechanisms of action. In contrast, the hERG + CaV group did not show significant differences from either the hERG or CaV groups, indicating that these two groups could not be completely distinguished. This finding suggests that for multi-channel inhibitors, the predominant inhibitory effect varies among compounds, and in some cases, the response patterns resemble those of single-channel inhibitors.

**Figure 6.**
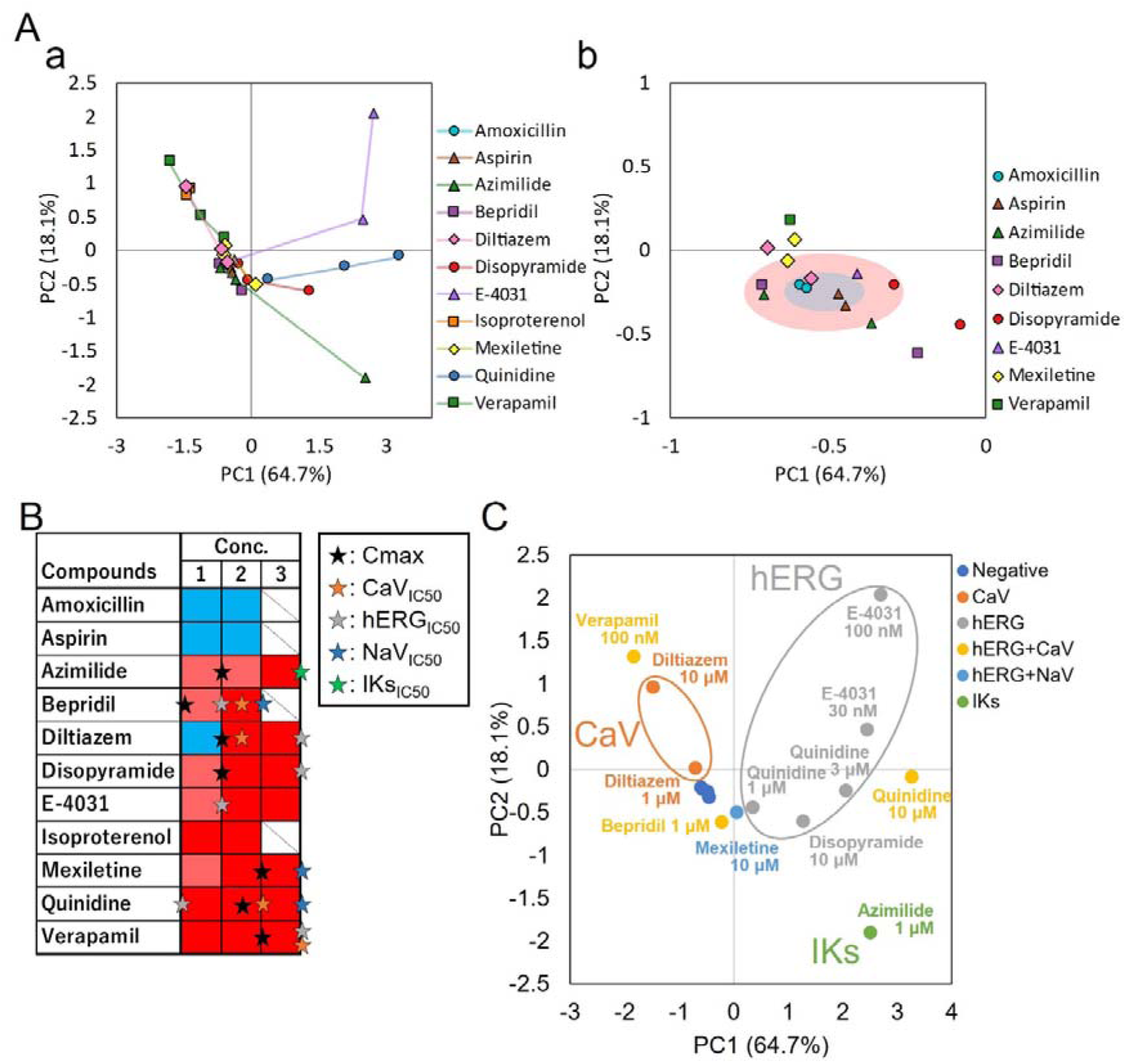
Detection of Cardiotoxicity and Classification of Mechanisms of Action Using PCA Analysis of Beat Parameters. (A) (a) PCA plot of compounds. (b) Enlarged view of the region near negative control compounds. (B) Toxicity classification based on the standard deviation range of negative control compounds. Stars indicate the IC₅₀ and Cmax of each ion channel.(C) Classification of mechanisms of action on the PCA plot based on IC₅₀ values.

**Table 3.** Principal component loadings. Beat rate, IBI, Area, and STV of Velocity, were identified to be an effective parameter set.

|  | **Principal component loadings** |  |  |  |
| --- | --- | --- | --- | --- |
|  | **Beat rate** | **IBI** | **Area** | **Velocity STV** |
| **PC1** | -0.49 | 0.54 | -0.51 | 0.46 |
| **PC2** | 0.60 | -00.033 | -0.32 | 0.66 |

**Table 4.** Summary of Cmax Free Concentrations and IC_50_ Values for Each Ion Channel.

| Compounds | Cmax Free (μM) | IC<sub>50</sub> (μM) |  |  |  |
| --- | --- | --- | --- | --- | --- |
|  |  | hERG | NaV | CaV | IKs |
| Azimilide | 0.07 | 5.2 | 19 | 17.8 | 0.7 |
| Bepridil | 0.032 | 0.16 | 2.3 | 1 | - |
| Diltiazem | 0.128 | 13.2 | 22.4 | 0.76 | - |
| Disopyramide | 0.7 | 14.4 | 168.4 | 1037 | - |
| E4031 | - | 0.014 | - | - | - |
| Mexiltetine | 2.5 | 62.2 | 38 | 125 | - |
| Quinidine | 3 | 0.72 | 14.6 | 6.4 | - |
| Verapamil | 0.045 | 0.25 | 32.5 | 0.2 | - |

**Table 5.** Statistical analysis of PCA using the effective parameter set (Beat rate, IBI, Area, and STV of Velocity)

|  | vs. Negative | vs. CaV | vs. hERG | vs. hERG + CaV | vs. hERG + NaV | vs. IKs |
| --- | --- | --- | --- | --- | --- | --- |
| **Negative** | - | p <0.001** | p <0.001** | p =0.009** | p =0.001** | p <0.001** |
| **CaV** | p <0.001** | - | p <0.001** | p =0.177 | p <0.001** | p <0.001** |
| **hERG** | p <0.001** | p <0.001** | - | p =0.326 | p =0.043* | p <0.001** |
| **hERG+CaV** | p <0.009** | p =0.177 | p =0.326 | - | p =0.044* | p <0.001** |
| **hERG + NaV** | p <0.001** | p <0.001** | p =0.043* | p =0.044* | - | p <0.001** |
| **IKs** | p <0.001** | p <0.001** | p <0.001** | p <0.001** | p <0.001** | - |

### Results 4. Detection of Chronic Cardiotoxicity Using Beat Parameters

To validate the chronic cardiotoxicity assessment using CMOS-MEA measurements, we conducted a chronic exposure experiment with doxorubicin. Doxorubicin (0.03 µM and 0.1 µM) and its solvent control, DMSO, were administered for 96 hours, and measurements were performed before exposure and every 24 hours thereafter. The histogram of active electrodes, delay map, and amplitude map obtained are shown in Figure 7A. Long-term exposure to DMSO did not result in any temporal changes in beat rate, beat area, or peak potential. In contrast, exposure to doxorubicin at 0.03 µM did not significantly alter the beat rate, but both beat area and peak potential significantly decreased after 24 hours. Furthermore, exposure to doxorubicin at 0.1 µM resulted in a substantial and time-dependent reduction in beat rate, beat area, and peak potential after 48 hours.

**Figure 7.**
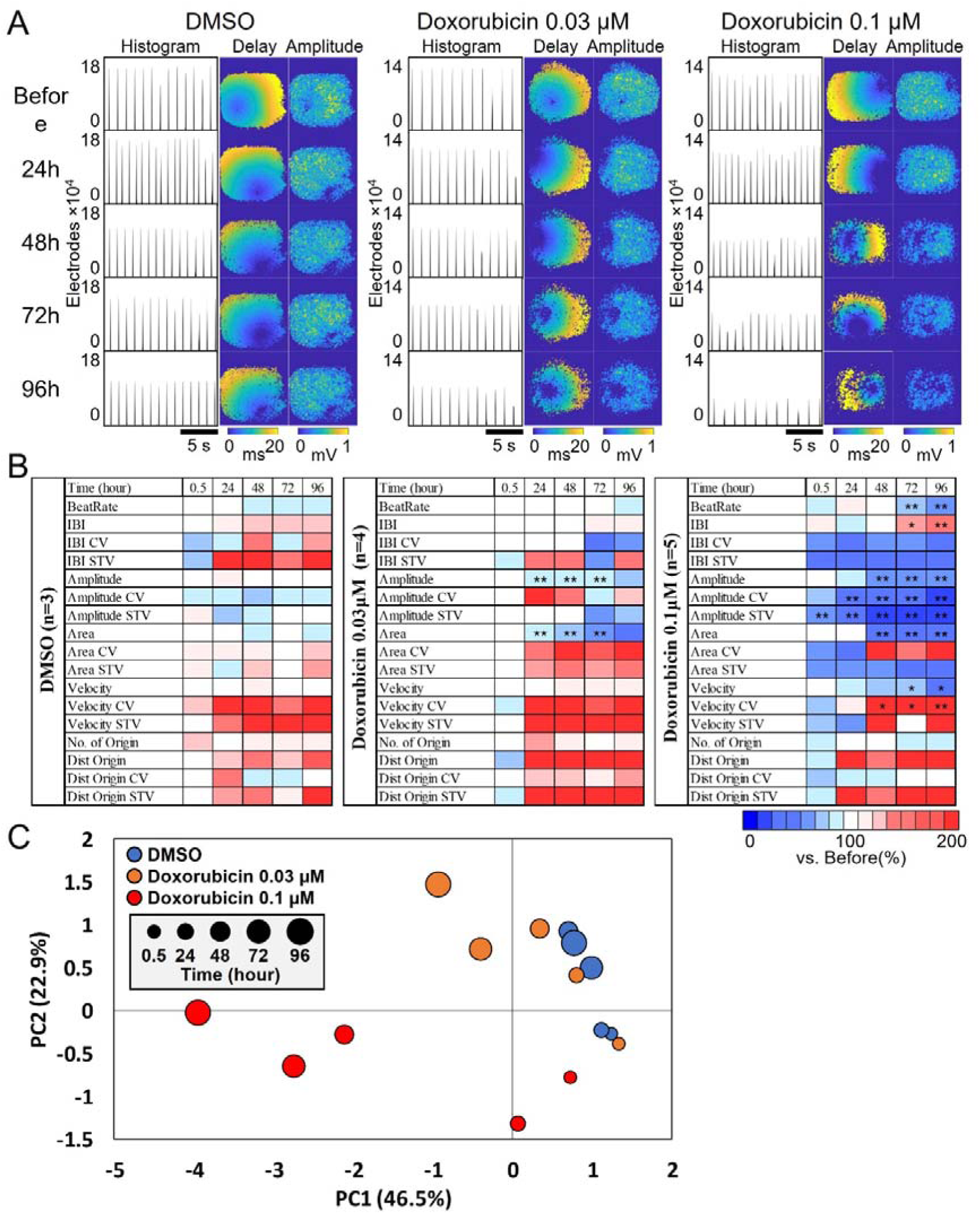
Detection of Chronic Cardiotoxicity Induced by Doxorubicin. (A) (a) Histogram of the number of active electrodes before and after chronic treatment (left), heatmap of peak timing for a single beat (middle), and heatmap of peak amplitude for a single beat (right). (B) Time-dependent heatmap of beat parameters after chronic treatment. (C) PCA plot after chronic treatment.

The detailed temporal changes in beat-related parameters are visualized as a heatmap in Figure 7B. During long-term exposure to DMSO, slight changes in the coefficient of variation (CV) and short-term variability (STV) were observed; however, these changes were not statistically significant. In contrast, exposure to doxorubicin at 0.03 µM led to a significant reduction in the newly developed parameters, beat area (Area) and peak potential (Amplitude), after 24 hours. At 0.1 µM, a reduction in the STV of peak potential was observed as early as 0.5 hours post-exposure, followed by a decrease in CV after 24 hours. Additionally, after 48 hours, an increase in the CV of conduction velocity was observed, along with the reduction of beat area and peak potential. Changes in conventional parameters, such as beat rate and IBI, which can be evaluated using traditional MEA, were detected after 72 hours of exposure. These results indicate that the newly developed parameters are more sensitive and can detect the effects of doxorubicin earlier than conventional parameters.

Furthermore, chronic exposure to doxorubicin affected multiple parameters, leading to variations in the interpretation of cardiotoxicity depending on the parameter of interest. To enable a comprehensive evaluation, we employed PCA to detect cardiotoxicity based on multiple parameter changes. Table 6 presents the parameters used for PCA and their loading values (cumulative contribution rate of PC1 and PC2: 69.4%). The results of PCA using IBI, Amplitude, Amplitude STV, Area, Velocity, and Velocity STV are shown in Figure 7C. Long-term exposure to DMSO resulted in minimal temporal transitions, with the plots remaining in the same region. In contrast, exposure to doxorubicin at 0.03 µM and 0.1 µM caused large temporal shifts, deviating from the DMSO cluster.

**Table 6.**
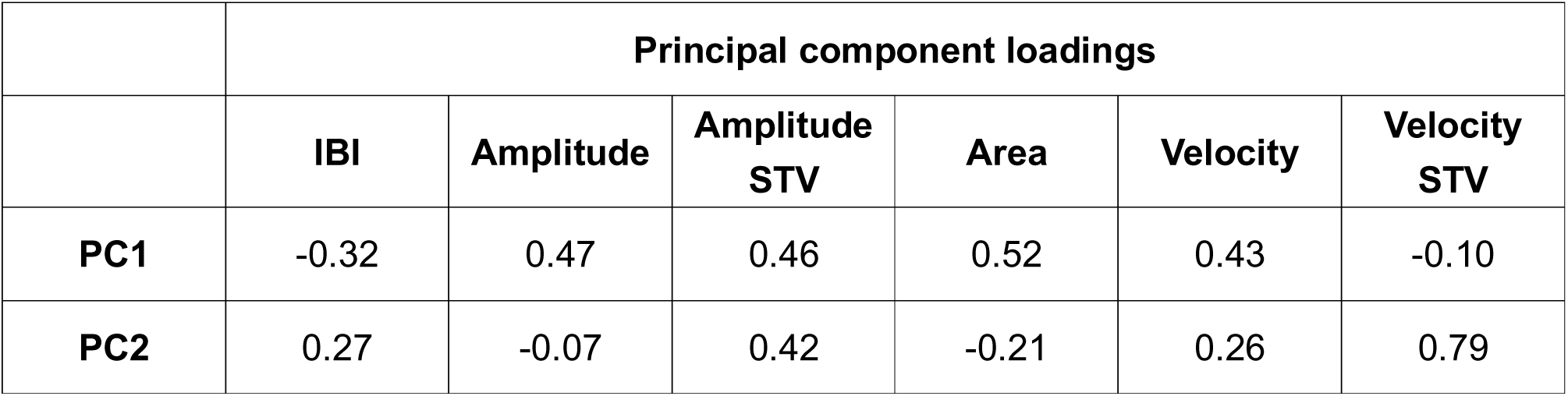
Principal component loadings. IBI, Amplitude, STV of Amplitude, Area, Velocity, and STV of Velocity, were identified to be an effective parameter set to remove the effect of time course of DMSO exposure.

|  | Principal component loadings |  |  |  |  |  |
| --- | --- | --- | --- | --- | --- | --- |
|  | IBI | Amplitude | Amplitude STV | Area | Velocity | Velocity STV |
| PC1 | -0.32 | 0.47 | 0.46 | 0.52 | 0.43 | -0.10 |
| PC2 | 0.27 | -0.07 | 0.42 | -0.21 | 0.26 | 0.79 |

To detect the chronic toxicity of doxorubicin based on the PCA plot, we classified the obtained plots into 15 groups based on concentration and time and performed MANOVA to assess changes relative to each time point of DMSO exposure. No significant differences were observed between the DMSO groups at any time point. For doxorubicin at 0.03 µM, significant differences were detected after 24 hours compared to 0.5 hours of DMSO exposure. For doxorubicin at 0.1 µM, significant differences were observed after 24 hours compared to all DMSO time points (Table 7).

**Table 7.** Statistical analysis of PCA using the effective parameter set (IBI, Amplitude, Amplitude STV, Area, Velocity, and Velocity STV) to eliminate the effect of time course of DMSO exposure.

| | DMSO | | | | | Doxorubicin 0.03 μM | | | | | Doxorubicin 0.1 μM | | | | |
| --- | --- | --- | --- | --- | --- | --- | --- | --- | --- | --- | --- | --- | --- | --- | --- |
|  | 0.5h | 24h | 48h | 72h | 96h | 0.5h | 24h | 48h | 72h | 96h | 0.5h | 24h | 48h | 72h | 96h |
| **DMSO 0.5 h** | · | p<br>=0.98<br>7 | p<br>=0.51<br>1 | p<br>=0.48<br>9 | p<br>=0.26<br>7 | p<br>=0.92<br>4 | p<br>=0.93<br>2 | p<br>=0.47<br>1 | p<br>=0.44<br>5 | p<br>=0.30<br>0 | p<br>=0.55<br>4 | p<br>=0.38<br>2 | p<br>=0.31<br>9 | p<br>=0.31<br>5 | p<br>=0.06<br>6 |
| **DMSO 24 h** | p<br>=0.98<br>7 | · | p<br>=0.68<br>4 | p<br>=0.80<br>2 | p<br>=0.39<br>7 | p<br>=0.04<br>7* | p<br>=0.64<br>1 | p<br>=0.81<br>3 | p<br>=0.96<br>4 | p<br>=0.82<br>8 | p<br>=0.03<br>3* | p<br>=0.01<br>4* | p<br>=0.01<br>6* | p<br>=0.02<br>9* | p<br>=0.00<br>1** |
| **DMSO 48 h** | p<br>=0.511 | p<br>=0.68<br>4 | · | p<br>=0.90<br>4 | p<br>=0.98<br>9 | p<br>=0.02<br>4* | p<br>=0.22<br>7 | p<br>=0.82<br>1 | p<br>=0.34<br>1 | p<br>=0.56<br>7 | **p**<br>**=0.002**<br>**\*\*** | **p**<br>**=0.00**<br>**1\*\*** | **p**<br>**=0.00**<br>**4\*\*** | **p**<br>**=0.00**<br>**6\*\*** | **p**<br>**=0.00**<br>**2\*\*** |
| **DMSO 72 h** | p<br>=0.48<br>9 | p<br>=0.80<br>2 | p<br>=0.90<br>4 | · | p<br>=0.91<br>7 | p<br>=0.03<br>0* | p<br>=0.13<br>5 | p<br>=0.37<br>3 | p<br>=0.18<br>7 | p<br>=0.21<br>6 | p<br>=0.00<br>8** | p<br>=0.00<br>2** | p<br>=0.00<br>8** | p<br>=0.01<br>8* | p<br>=0.00<br>1** |
| **DMSO 96 h** | p<br>=0.26<br>7 | p<br>=0.39<br>7 | p<br>=0.98<br>9 | p<br>=0.91<br>7 | · | p<br>=0.01<br>7* | p<br>=0.06<br>0 | p<br>=0.16<br>8 | p<br>=0.07<br>6 | p<br>=0.08<br>0 | p<br>=0.00<br>4** | p<br><0.00<br>1** | p<br>=0.00<br>5** | p<br>=0.01<br>2* | p<br><0.00<br>1** |

These results demonstrate that the novel chronic toxicity assessment method, which integrates CMOS-MEA-based newly developed parameters with PCA, enables highly sensitive detection of doxorubicin-induced cardiotoxicity at lower concentrations and shorter exposure durations.

## Discussion

FPI using CMOS-MEA offers high-resolution measurements capable of simultaneously recording feild potentials from a single cell across multiple electrodes. This allows for the analysis of the electrophysiological activity of entire human iPSC-derived cardiomyocyte networks with high spatial and temporal resolution (Figure 1). The novel parameters derived for the first time through FPI, including the number of excitation origins, propagation velocity, and propagation area, have been demonstrated to serve as effective endpoints for explaining drug mechanisms.

Specifically, in addition to an increase in beating frequency, an increase in the number of excitation origins was observed with Isoproterenol. This suggests that Isoproterenol, a β-adrenergic agonist, captures the characteristic arrhythmogenic effects in cardiomyocytes, including the induction of early afterdepolarizations (EADs) and/or delayed afterdepolarizations (DADs) (Figure 2)^35, 36^. With Mexiletine, a decrease in beating frequency was observed at 10 μM, indicating that the bradycardic effect of Mexiletine was also detected in MEA measurements. Furthermore, a reduction in propagation velocity was observed, suggesting that the restricted influx of sodium ions delays the propagation of ionic excitation between cells (Figure 3). For E-4031, a decrease in propagation area and the detection of EAD-like waveforms were observed, revealing conduction abnormalities that reflect the arrhythmia risk associated with hERG channel inhibition (Figure 4). As seen with E-4031, when partial beats occur, conventional MEA measurements with fewer electrodes may struggle to detect these beats. The novel parameters derived from FPI serve as effective endpoints that reflect mechanisms of action and have the potential to establish FPI as a next-generation myocardial MEA assay for predicting drug-induced cardiotoxicity risks. Furthermore, in cases where multiple beating patterns occur, conventional MEA calculates IBIs based on the peak timing detected by a single electrode. As a result, it fails to accurately detect IBIs and their coefficient of variation at the cellular population level. In contrast, since FPI records the activity of the entire cardiomyocyte population, it offers the advantage of providing more precise values for fundamental parameters such as beat rate and IBI. Amplitude-related parameters also showed significant changes in the acute response of many compounds (Figure 5), and doxorubicin in chronic treatment assays also exhibited prominent changes. This indicates that the amplitude calculated by the UHD-CMOS-MEA system, derived from over 100,000 active electrodes capable of recording the activity of the entire cell population, exhibits higher sensitivity in detection. Comparison of cardiotoxicity detection using novel FPI endpoints and conventional methods revealed that changes induced by Verapamil and Mexiletine were detected at lower concentrations than in traditional assays. Furthermore, FPI enabled the detection of changes induced by Bepridil, which were difficult to capture using conventional MEA measurements (Table 8), demonstrating its potential contribution to advancing in vitro cardiotoxicity assessment.

**Table 8.**
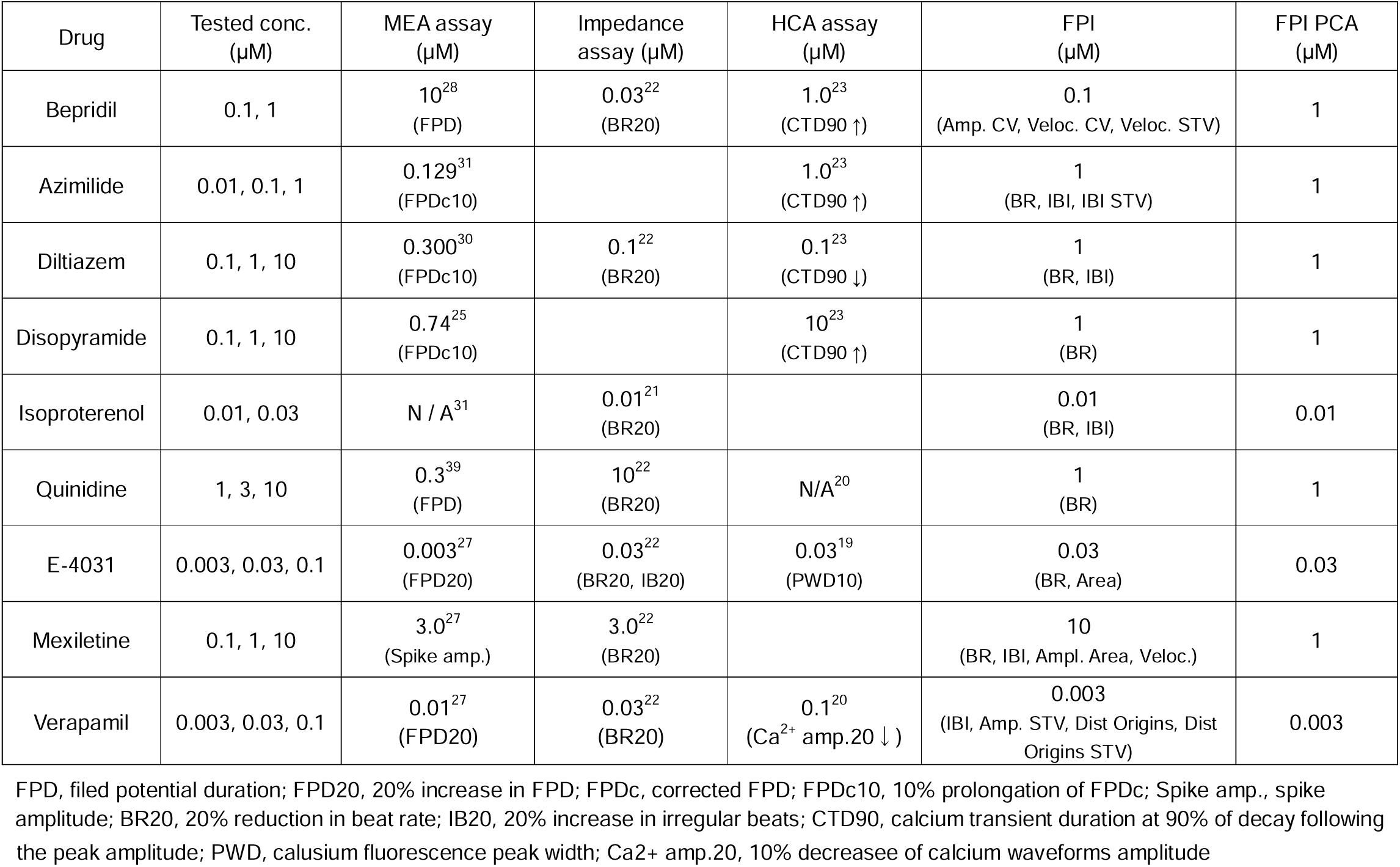
Comparison with Cardiotoxicity assay benchmark.

The results of PCA using the novel endpoints, Area and Velocity STV, along with Beat rate and IBI, demonstrated the ability to clearly capture concentration-dependent changes and evaluate complex toxic effects, including Bepridil, which was difficult to detect using conventional MEA measurements and is classified as an outlier compound in CiPA (Figure 5A). Bepridil is a hERG blocker with a tendency to prolong repolarization; however, due to its additional properties as a Ca²⁺ and Na⁺ channel antagonist, the extent of repolarization prolongation tends to be counterbalanced, making cardiotoxicity detection challenging^12^. The multivariate analysis incorporating novel endpoints, combined with an evaluation method based on the standard deviation range of negative controls, suggests its effectiveness as a cardiotoxicity assessment method (Figure 5A). Compounds acting on multiple ion channels exert their combined effects on cells and tissues. Conventional IC₅₀-based toxicity risk assessments, particularly those focused on hERG inhibition, have faced challenges due to discrepancies between in vitro assessments and clinically toxic doses^37, 38^. hiPSC-derived cardiomyocyte assays, which inherently possess multiple ion channels, enable simultaneous multi-channel evaluation^14^. The FPI technology and multivariate analysis incorporating novel endpoints enable toxicity assessment that accounts for multichannel effects, contributing to improved accuracy in clinical toxicity prediction.

Furthermore, in the classification of drug mechanisms of action using PCA, compound groups with a single mechanism, such as those acting on hERG, Na, and Ca channels, were significantly separated, demonstrating that mechanism-based prediction is feasible (Table 5). Additionally, the classification of compounds acting on multiple channels was examined. As a result, Diltiazem 10 μM (hERG: 76%, NaV: 45%, CaV: 1316%, with each value representing the ratio relative to IC_50_) was plotted near compounds acting on CaV channels (Figure 6C). Similarly, Quinidine 10 μM (hERG: 1389%, NaV: 68%, CaV: 156%, with each value representing the ratio relative to IC_50_) was plotted near compounds acting on hERG channels (Figure 6C). The coordinates of the PCA plot reflected differences in IC_50_ values across channels for each compound, demonstrating that even when a compound acts on multiple channels, its predominant mechanism of action can still be evaluated.

Moreover, these findings suggest that the combination of FPI and PCA analyses contributes not only to cardiotoxicity risk assessment but also to the elucidation of mechanisms of action (Fig. 6). Predicting mechanisms of action can facilitate lead compound optimization in the early stages of drug discovery and is also expected to be useful for predicting off-target effects.

In the chronic toxicity assessment of doxorubicin, we detected significant signs of cardiotoxicity at a low concentration (0.03 µM) and within a short duration (24 hours). Specifically, while no clear changes were observed in beating frequency, early reductions in propagation area and peak potential were detected, successfully capturing the early signs of electrical desynchronization and remodeling in cardiomyocytes (Figure 7A, 7B). Furthermore, at a high concentration (0.1 µM), a decrease in conduction velocity and an increase in the coefficient of variation (CV, STV) were observed after 48 hours, indicating a significant impact on the propagation characteristics of cardiomyocytes. A previous study using mouse cardiomyocytes reported that treatment with high concentrations of doxorubicin (1 µM) led to a reduction in contractile force, along with impaired Ca²⁺ uptake and release capacity, affecting Ca² ⁺ handling^40^. Additionally, studies using rat cardiomyocytes have shown that doxorubicin-induced abnormalities in Ca² ⁺ handling trigger apoptosis in cardiomyocytes, leading to reduced contractility^41, 42^. The effects of doxorubicin on the peak field potential of cardiomyocytes, as captured by the CMOS-MEA, may reflect abnormalities in Ca handling at the level of individual cardiomyocytes. Additionally, the effects on propagation area and conduction velocity may indicate reduced electrical synchrony between cells due to Ca handling abnormalities. A previous study using engineered heart tissues (EHTs) derived from human iPS cardiomyocytes reported a decrease in contractile force following 48-hour treatment with 1 µM doxorubicin, while no reduction in contractile force was observed at lower concentrations of 125 and 500 nM^43^. Furthermore, in a study utilizing deep learning-based cardiotoxicity assessment of human iPSC-derived cardiomyocytes in monolayer cultures, structural changes in cells were detected from images before the decrease in contractile force following 24- and 48-hour doxorubicin treatment. However, the detection threshold was above 0.3 µM, and no changes were observed at 0.1 µM^44^. In comparison to these previous studies, our findings demonstrate that CMOS-MEA enabled the detection of cardiotoxicity signs at an low concentration (0.03 µM) and in a short period (24 hours). This suggests that CMOS-MEA has a higher sensitivity than conventional methods, as it can assess cardiomyocyte function at the network level, locally, and even at the single-cell level, beyond merely evaluating cell morphology (adhesion) or viability. These results indicate that CMOS-MEA possesses the capability to detect the early stages of toxicity progression more efficiently than conventional assessment approaches.

The novel endpoints derived from FPI using the UHD-CMOS-MEA system, which provides high spatial and temporal resolution—such as the number of excitation origins, conduction velocity, and propagation area—combined with multivariate analysis for cardiotoxicity assessment, represent a next-generation in vitro cardiotoxicity prediction platform. This platform is expected to enable not only the risk assessment of cardiotoxicity across a broad range of drug candidates but also the elucidation of mechanisms of action and the prediction of off-target effects.

## Materials and Methods

### Chip Preparation

MEA chips were pretreated with a Tergazyme solution prepared by dissolving 0.5 g of Tergazyme in 50 mL of sterile water at 60°C. Each well was filled with 3 mL of the solution and incubated for 1 hour at room temperature. The chips were then washed four times with 3 mL of distilled water (DW). Next, 3 mL of 70% ethanol was added to each well for 30 seconds, followed by four additional DW washes. After the final wash, the chips were placed in a biosafety cabinet and exposed to UV light for 30 minutes to ensure sterilization while allowing them to air-dry. The dried chips were then coated with a Type-C collagen solution (diluted 10-fold in 0.02 N acetic acid) and incubated overnight at 4°C. After removing the collagen solution, the chips were air-dried at room temperature for 2 hours. To enhance cell adhesion, a fibronectin solution (50 µg/mL, prepared by diluting a 1 mg/mL stock solution 20-fold in PBS) was added directly to the electrode area (15 µL per chip) and incubated at 37°C for 1 hour.

### Cell Culture and Plating

Cryopreserved iCell Cardiomyocytes (FUJIFILM Cellular Dynamics, Inc.) were handled according to the supplier’s instructions. Cells were thawed in a 37°C water bath for approximately 3 minutes and resuspended in iCell Thawing Medium to prepare a cell suspension at a final concentration of 1×10[ cells/mL. A total of 10 µL of the cell suspension (100,000 cells per chip) was seeded directly onto the fibronectin-coated electrode area of each chip. Chips were incubated at 37°C for 1 hour to allow cell attachment, after which 1 mL of maintenance medium was gently added. The culture medium was replaced 24 hours after seeding, with half the volume exchanged. Thereafter, medium exchanges were performed twice weekly by replacing half the medium volume.

### Immunocytochemistry

Immunocytochemistry was performed on human iPSC-derived cardiomyocytes cultured on CMOS-MEA chips immediately after pharmacological experiments. Stain Perfect Kit A (#SP-A-1000, Immusmol) was used for the staining procedure. CMOS-MEA chips were first gently washed with Wash Solution 1, followed by fixation with Fixation Solution at room temperature for 10 minutes. After fixation, the chips were washed three times with Wash Solution 1 and permeabilized using Permeabilization Solution at room temperature for 10 minutes. The chips were then washed again three times with Wash Solution 1 and stabilized with Stabilization Solution for 10 minutes at room temperature. Saturation Solution was subsequently applied and incubated for 30 minutes at room temperature to block non-specific binding. For primary antibody staining, MLC-2A antibody (Synaptic Systems, 311011) and MLC-2V antibody (ProteinTech, 10906-1-AP) were diluted 1:200 in the appropriate buffer and applied to the chips. The chips were incubated overnight at 4°C. The next day, the chips were washed three times with Wash Solution 2, and secondary antibodies, Goat anti-Mouse IgG (H+L) Cross-Adsorbed Secondary Antibody, Alexa Fluor™ 488 (Invitrogen, A-11001), and Goat anti-Rabbit IgG (H+L) Highly Cross-Adsorbed Secondary Antibody, Alexa Fluor™ 546 (Invitrogen, A-11035), were diluted 1:1000 and applied at room temperature for 30 minutes. After washing three times with Wash Solution 2, nuclei were stained with Hoechst 33258 solution (DOJINDO, 34307961) diluted 1:1000, and incubated for 10 minutes. The chips were washed three more times with Wash Solution 3, and the samples were mounted using ProLong Glass Antifade Mountant (Invitrogen, P36984). Images were captured using a confocal laser scanning microscope (Nikon, AX/AXR with NSPARK) equipped with a 20× objective lens.

### Pharmacological Tests

Pharmacological tests were conducted on human iPSC-derived cardiomyocytes after 1 week of culture. Details of the test compounds are summarized in Table 9. All drugs were dissolved in 0.1% DMSO. Spontaneous firing was recorded for 2 minutes at each concentration, with compounds administered cumulatively. During recording and drug treatment, cultures were maintained at 37 °C in a 5% CO₂ atmosphere.

**Table 9.** Details of test compounds.

| Compound | Class | Conc. (μM) | Source | CAS RN® |
| --- | --- | --- | --- | --- |
| Amoxicillin | β-lactam antibiotic | 10, 100 | A2099, Tokyo Chemical Industry Co. | 61336-70-7 |
| Aspirin | Analgesic, anti-pyretic | 10, 100 | A2093, Sigma-Aldrich | 50-78-2 |
| Azimilide | Class III antiarrhythmic | 0.01, 0.1, 1 | 6318, Tocris Bioscience | 149888-94-8 |
| Bepridil | Anti-anginal | 0.1, 1 | B5016, Sigma-Aldrich | 68099-86-5 |
| Diltiazem | Class IV antiarrhythmic | 0.1, 1, 10 | 047-20311, Fujifilm Wako Pure Chemical Co. | 33286-22-5 |
| Disopyramide | Class Ia antiarrhythmic | 0.1, 1, 10 | S5490, Selleck Chemicals | 3737-09-5 |
| E-4031 | Class III anti-arrhythmic (research purposes only) | 0.003, 0.03, 0.1 | 059-08451, Fujifilm Wako Pure Chemical Co. | N/A-05-0845-1 |
| Isoproterenol | Class II anti-arrhythmic | 0.01, 0.03 | I0260, Tokyo Chemical Industry Co. | 51-30-9 |
| Mexiletine | Class Ib anti-arrhythmic | 0.1, 1, 10 | 132-17581, Fujifilm Wako Pure Chemical Co. | 5370-01-4 |
| Quinidine | Class Ia antiarrhythmic | 1, 3, 10 | 176-00111, Fujifilm Wako Pure Chemical Co. | 56-54-2 |
| Verapamil | Class IV anti-arrhythmic | 0.003, 0.03, 0.1 | 222-00781, Fujifilm Wako Pure Chemical Co. | 152-11-4 |
| DMSO | Vehicle | 0.1% | D2650, Sigma-Aldrich | 67-68-5 |
| Doxorubicin | Broad-spectrum anti-tumor | 0.03, 0.1 | D4193, Tokyo Chemical Industry Co. | 25316-40-9 |

### Extracellular Recording

Spontaneous extracellular field potentials were recorded using a high-density CMOS-MEA system (Sony Semiconductor Solutions) at 37 °C in a 5% CO[ atmosphere. Measurements were performed with 236,880 electrodes at a sampling rate of 2 kHz, and recording sessions lasted for 2 minutes per condition. Data acquisition was controlled by dedicated software (Sony Semiconductor Solutions), and all raw data were stored on a PC for subsequent analysis.

### Spike Detection

Spikes at each electrode of hiPSC-derived cardiomyocyte samples were detected using a voltage threshold of ±100 µV. Spike detection was performed using BinViewer software (SCREEN Holdings Co., Ltd., Japan).

### Beat Detection

Beat detection was performed using a custom program developed in MATLAB (MathWorks, Natick, MA). Spike detection results were used to generate firing count histograms with a bin size of 0.5 ms. The timing and propagation duration of pulses across electrodes were identified through peak detection in these histograms. For each electrode, the maximum amplitude and its corresponding peak timing were calculated, enabling the generation of amplitude and peak timing heat maps.

### Principal component analysis

A matrix comprising 17 analytical parameters was constructed for PCA. The MATLAB PCA function was used to analyze 65,518 parameter sets, allowing the selection of 2 to 8 parameters from the 17 available. A one-way multivariate analysis of variance (MANOVA) was performed on the PCA results to assess significant differences in the first two principal components among the negative control compounds. The parameter set with no significant differences among negative control compounds was extracted for subsequent analysis.

### Statistical Analysis

One-way ANOVA was performed for each of the 17 calculated beat parameters to assess whether the values at each concentration differed significantly from the pre-treatment baseline. For mechanism-of-action (MoA) classification and the cardiotoxicity assessment of doxorubicin, a one-way multivariate ANOVA (MANOVA) was employed, using PC1 and PC2 scores as dependent variables to evaluate significant differences between drug mechanism groups. Statistical significance was determined at p<0.05.

### Cardiotoxicity assessment

The optimized analytical parameters of each testing compound were compared with the SD of negative compounds to predict the cardiotoxicity potential on a relatively quantitative scale. For example, a low risk was predicted for values below the SD range, a medium risk for those with 2 SD, and a high risk for those with >2 SD.

## Acknowledgments

This study was supported by the grant of Japan Agency for Medical Research and Development (AMED) Grant Number 23be1004203h0002 and 24be1004203h0003. This study was also supported by the UHD-CMOS-MEA system with Sony semiconductor solutions Inc.

## Author contributions

Naoki Matsuda and Nami Nagafuku: These authors contributed equally to this work. All authors designed experiments; N.Nagafuku conducted experiments; N.Matsuda, K.Matsuda, and Y.Ishibashi analyzed the data; I.Suzuki, N.Nagafuku, N.Matsuda, and Y.Ishibashi wrote the manuscript; T. Taniguchi, Y. Matsushita, N. Miyamoto, and T. Yoshinaga investigation & review; I.Suzuki supervised the project.

## References

1. Cook D, et al. Lessons learned from the fate of AstraZeneca’s drug pipeline: a five-dimensional framework. Nature reviews Drug discovery 13, 419–431 (2014).

2. Fung M, Thornton A, Mybeck K, Wu JH-h, Hornbuckle K, Muniz E. Evaluation of the characteristics of safety withdrawal of prescription drugs from worldwide pharmaceutical markets-1960 to 1999. Drug Information Journal 35, 293–317 (2001).

3. Mamoshina P, Rodriguez B, Bueno-Orovio A. Toward a broader view of mechanisms of drug cardiotoxicity. Cell Reports Medicine 2, (2021).

4. Onakpoya IJ, Heneghan CJ, Aronson JK. Post-marketing withdrawal of 462 medicinal products because of adverse drug reactions: a systematic review of the world literature. BMC medicine 14, 1–11 (2016).

5. Stockbridge N, Morganroth J, Shah RR, Garnett C. Dealing with global safety issues: was the response to QT-liability of non-cardiac drugs well coordinated? Drug safety 36, 167–182 (2013).

6. Committee IS. The nonclinical evaluation of the potential for delayed ventricular repolarization (QT interval prolongation) by human pharmaceuticals (S7B). In: The international conference on harmonization of technical requirements for registration of pharmaceuticals for human use (ICH). The Guideline was recommended for adoption at Step 4 of the ICH process in May 2005) (2005).

7. Gintant G. An evaluation of hERG current assay performance: Translating preclinical safety studies to clinical QT prolongation. Pharmacology & therapeutics 129, 109–119 (2011).

8. Colatsky T, et al. The comprehensive in vitro proarrhythmia assay (CiPA) initiative—update on progress. Journal of pharmacological and toxicological methods 81, 15–20 (2016).

9. Sager PT, Gintant G, Turner JR, Pettit S, Stockbridge N. Rechanneling the cardiac proarrhythmia safety paradigm: a meeting report from the Cardiac Safety Research Consortium. American heart journal 167, 292–300 (2014).

10. Yang X, Papoian T. Moving beyond the comprehensive in vitro proarrhythmia assay: Use of human-induced pluripotent stem cell-derived cardiomyocytes to assess contractile effects associated with drug - induced structural cardiotoxicity. Journal of Applied Toxicology 38, 1166–1176 (2018).

11. Li Z, et al. Assessment of an in silico mechanistic model for proarrhythmia risk prediction under the ci pa initiative. Clinical Pharmacology & Therapeutics 105, 466–475 (2019).

12. Blinova K, et al. International multisite study of human-induced pluripotent stem cell-derived cardiomyocytes for drug proarrhythmic potential assessment. Cell reports 24, 3582–3592 (2018).

13. Kanda Y, Yamazaki D, Osada T, Yoshinaga T, Sawada K. Development of torsadogenic risk assessment using human induced pluripotent stem cell-derived cardiomyocytes: Japan iPS Cardiac Safety Assessment (JiCSA) update. Journal of pharmacological sciences 138, 233–239 (2018).

14. Lee TY, Coles JG, Maynes JT. iPSC-cardiomyocytes in the preclinical prediction of candidate pharmaceutical toxicity. Frontiers in Pharmacology 15, 1308217 (2024).

15. Blinova K, et al. Comprehensive translational assessment of human-induced pluripotent stem cell derived cardiomyocytes for evaluating drug-induced arrhythmias. Toxicological Sciences 155, 234–247 (2017).

16. Gibson JK, Yue Y, Bronson J, Palmer C, Numann R. Human stem cell-derived cardiomyocytes detect drug-mediated changes in action potentials and ion currents. Journal of pharmacological and toxicological methods 70, 255–267 (2014).

17. Hoekstra M, Mummery CL, Wilde AA, Bezzina CR, Verkerk AO. Induced pluripotent stem cell derived cardiomyocytes as models for cardiac arrhythmias. Frontiers in physiology 3, 346 (2012).

18. Ma J, et al. High purity human-induced pluripotent stem cell-derived cardiomyocytes: electrophysiological properties of action potentials and ionic currents. American Journal of Physiology-Heart and Circulatory Physiology 301, H2006–H2017 (2011).

19. Aikawa N. The utility of human iPS cell-derived cardiomyocytes in predicting the clinical risk of drugs that display discordance of cardiotoxicity by species. Fundamental Toxicological Sciences 4, 127–136 (2017).

20. Dempsey GT, et al. Cardiotoxicity screening with simultaneous optogenetic pacing, voltage imaging and calcium imaging. Journal of pharmacological and toxicological methods 81, 240–250 (2016).

21. Doherty KR, Talbert DR, Trusk PB, Moran DM, Shell SA, Bacus S. Structural and functional screening in human induced-pluripotent stem cell-derived cardiomyocytes accurately identifies cardiotoxicity of multiple drug types. Toxicology and Applied Pharmacology 285, 51–60 (2015).

22. Guo L, Coyle L, Abrams RMC, Kemper R, Chiao ET, Kolaja KL. Refining the Human iPSC-Cardiomyocyte Arrhythmic Risk Assessment Model. Toxicological Sciences 136, 581–594 (2013).

23. Lu HR, et al. Assessing drug-induced long QT and proarrhythmic risk using human stem-cell-derived cardiomyocytes in a Ca2+ imaging assay: evaluation of 28 CiPA compounds at three test sites. Toxicological Sciences 170, 345–356 (2019).

24. Pointon A, Abi-Gerges N, Cross MJ, Sidaway JE. Phenotypic profiling of structural cardiotoxins in vitro reveals dependency on multiple mechanisms of toxicity. toxicological sciences 132, 317–326 (2013).

25. Ando H, et al. A new paradigm for drug-induced torsadogenic risk assessment using human iPS cell-derived cardiomyocytes. Journal of pharmacological and toxicological methods 84, 111–127 (2017).

26. Clements M, Millar V, Williams AS, Kalinka S. Bridging functional and structural cardiotoxicity assays using human embryonic stem cell-derived cardiomyocytes for a more comprehensive risk assessment. Toxicological sciences 148, 241–260 (2015).

27. Clements M, Thomas N. High-throughput multi-parameter profiling of electrophysiological drug effects in human embryonic stem cell derived cardiomyocytes using multi-electrode arrays. Toxicological Sciences 140, 445–461 (2014).

28. Kaneko T, et al. On-chip in vitro cell-network pre-clinical cardiac toxicity using spatiotemporal human cardiomyocyte measurement on a chip. Scientific reports 4, 4670 (2014).

29. Millard D, et al. Cross-site reliability of human induced pluripotent stem cell-derived cardiomyocyte based safety assays using microelectrode arrays: results from a blinded CiPA pilot study. Toxicological Sciences 164, 550–562 (2018).

30. Nozaki Y, Honda Y, Tsujimoto S, Watanabe H, Kunimatsu T, Funabashi H. Availability of human induced pluripotent stem cell-derived cardiomyocytes in assessment of drug potential for QT prolongation. Toxicology and applied pharmacology 278, 72–77 (2014).

31. Nozaki Y, et al. CSAHi study-2: validation of multi-electrode array systems (MEA60/2100) for prediction of drug-induced proarrhythmia using human iPS cell-derived cardiomyocytes: assessment of reference compounds and comparison with non-clinical studies and clinical information. Regulatory Toxicology and Pharmacology 88, 238–251 (2017).

32. Emery BA, et al. Large-scale multimodal recordings on a high-density neurochip: olfactory bulb and hippocampal networks. In: 2022 44th Annual International Conference of the IEEE Engineering in Medicine & Biology Society (EMBC)). IEEE (2022).

33. Suzuki I, et al. Large-Area Field Potential Imaging Having Single Neuron Resolution Using 236 880 Electrodes CMOS-MEA Technology. Advanced Science 10, 2207732 (2023).

34. Yuan X, Hierlemann A, Frey U. Extracellular recording of entire neural networks using a dual-mode microelectrode array with 19 584 electrodes and high SNR. IEEE journal of solid-state circuits 56, 2466–2475 (2021).

35. Paci M, Penttinen K, Pekkanen-Mattila M, Koivumäki JT. Arrhythmia mechanisms in human induced pluripotent stem cell–derived cardiomyocytes. Journal of Cardiovascular Pharmacology 77, 300–316 (2021).

36. Zeng J, Rudy Y. Early afterdepolarizations in cardiac myocytes: mechanism and rate dependence. Biophysical journal 68, 949–964 (1995).

37. Li Z, et al. Improving the In Silico Assessment of Proarrhythmia Risk by Combining hERG (Human Ether-à-go-go-Related Gene) Channel–Drug Binding Kinetics and Multichannel Pharmacology. Circulation: Arrhythmia and Electrophysiology 10, e004628 (2017).

38. Romero L, et al. In Silico QT and APD Prolongation Assay for Early Screening of Drug-Induced Proarrhythmic Risk. Journal of Chemical Information and Modeling 58, 867–878 (2018).

39. Harris K, Aylott M, Cui Y, Louttit JB, McMahon NC, Sridhar A. Comparison of Electrophysiological Data From Human-Induced Pluripotent Stem Cell–Derived Cardiomyocytes to Functional Preclinical Safety Assays. Toxicological Sciences 134, 412–426 (2013).

40. Lin R, et al. Empagliflozin attenuates doxorubicin-impaired cardiac contractility by suppressing reactive oxygen species in isolated myocytes. Molecular and Cellular Biochemistry 479, 2105–2118 (2024).

41. Ikeda S, et al. Blockade of L-type Ca2+ channel attenuates doxorubicin-induced cardiomyopathy via suppression of CaMKII-NF-κB pathway. Scientific Reports 9, 9850 (2019).

42. Sag CM, Köhler AC, Anderson ME, Backs J, Maier LS. CaMKII-dependent SR Ca leak contributes to doxorubicin-induced impaired Ca handling in isolated cardiac myocytes. Journal of Molecular and Cellular Cardiology 51, 749–759 (2011).

43. Arefin A, Mendoza M, Dame K, Garcia MI, Strauss DG, Ribeiro AJS. Reproducibility of drug-induced effects on the contractility of an engineered heart tissue derived from human pluripotent stem cells. Frontiers in Pharmacology 14, (2023).

44. Maddah M, Mandegar MA, Dame K, Grafton F, Loewke K, Ribeiro AJS. Quantifying drug-induced structural toxicity in hepatocytes and cardiomyocytes derived from hiPSCs using a deep learning method. Journal of Pharmacological and Toxicological Methods 105, 106895 (2020).

